# Deciphering the mechanism of inhibition of SERCA1a by sarcolipin using molecular simulations

**DOI:** 10.1101/2019.12.17.879825

**Authors:** T. Barbot, V. Beswick, C. Montigny, E. Quiniou, N. Jamin, L. Mouawad

**Affiliations:** Institut Curie; CEA Saclay; CEA

**Keywords:** Normal Modes Analysis, Molecular Modeling, Molecular Simulations, Calcium ATPase, SERCA1a, Sarcolipin

## Abstract

SERCA1a is an ATPase calcium pump that transports Ca^2+^ from the cytoplasm to the sarco/endoplasmic reticulum lumen. Sarcolipin (SLN), a transmembrane peptide, regulates the activity of SERCA1a by decreasing its Ca^2+^ transport rate, but its mechanism of action is still not well understood. To decipher this mechanism, we have performed normal modes analysis in the all-atom model, with the SERCA1a-SLN complex or the isolated SERCA1a embedded in an explicit membrane. The comparison of the results allowed us to provide an explanation for the action of SLN that is in good agreement with experimental observations. In our analyses, the presence of SLN locally perturbs the TM6 transmembrane helix and as a consequence modifies the position of D800, one of the key metal-chelating residues. Additionally, it reduces the flexibility of the gating residues, V304 and E309 in TM4, at the entrance of the Ca^2+^ binding sites, which would decrease the affinity for Ca^2+^. Unexpectedly, SLN has also an effect on the ATP binding site more than 35 Å away, due to the straightening of TM5, a long helix considered as the spine of the protein. The straightening of TM5 modifies the structure of the P-N linker that sits above it, and which comprises the ^351^DKTG^354^ conserved motif, resulting in an increase of the distance between ATP and the phosphorylation site. As a consequence, the turn-over rate could be affected. All this gives SERCA1a the propensity to go toward a Ca^2+^-deprived E2-like state in the presence of SLN and toward a Ca^2+^ high-affinity E1-like state in the absence of SLN, although the SERCA1a-SLN complex was crystallized in an E1-like state. In addition to a general mechanism of inhibition of SERCA1a regulatory peptides, this study also provides an insight in the conformational transition between the E2 and E1 states.

**Statement of Significance:** The role of sarco/endoplasmic reticulum calcium ATPase in muscle relaxation is essential. Impairment of its function may result in either cardiac diseases, or myopathies, and also thermogenesis defects. Inhibition of the ATPase by regulatory peptide such as sarcolipin remains unclear. The structure of the ATPase in complex with this peptide was studied by all-atom normal modes analysis, an in silico technique which allows us to decipher the mechanism of inhibition of calcium transport by sarcolipin at a molecular level. Our results open the way to understanding the impact of in vivo misregulation of the ATPase activity by sarcolipin. Development of tools enhancing or preventing interaction between the ATPase and its regulatory peptide could be considered as new therapeutic approaches.

## Introduction

The Sarco/Endoplasmic Reticulum Ca^2+^ ATPase (SERCA) is a transmembrane protein that transports Ca^2+^ from the cytoplasm to the Sarco/Endoplasm, using ATP as an energy source. In the skeletal muscles, SERCA1a, an isoform of the SERCA1 subfamily, is responsible for the muscle relaxation by restocking into the sarcoplasmic reticulum (SR) lumen the calcium that was formerly released in the cytosol during muscle contraction. This restocking results in a rapid lowering of the cytosolic Ca^2+^ concentration, which decreases the myosin/actin filaments activity, allowing for muscle relaxation (1, 2). SERCA1a is a 110 kDa protein composed of 10 transmembrane helices (TM1-10) and a large cytoplasmic headpiece that consists of three domains: the Nucleotide-binding (N) domain, which binds an ATP molecule, the Phosphorylation (P) domain, which contains the autophosphorylation site and the Actuator (A) domain, which is responsible for the dephosphorylation of the protein (Figure 1A) (3, 4).

**Figure 1.**
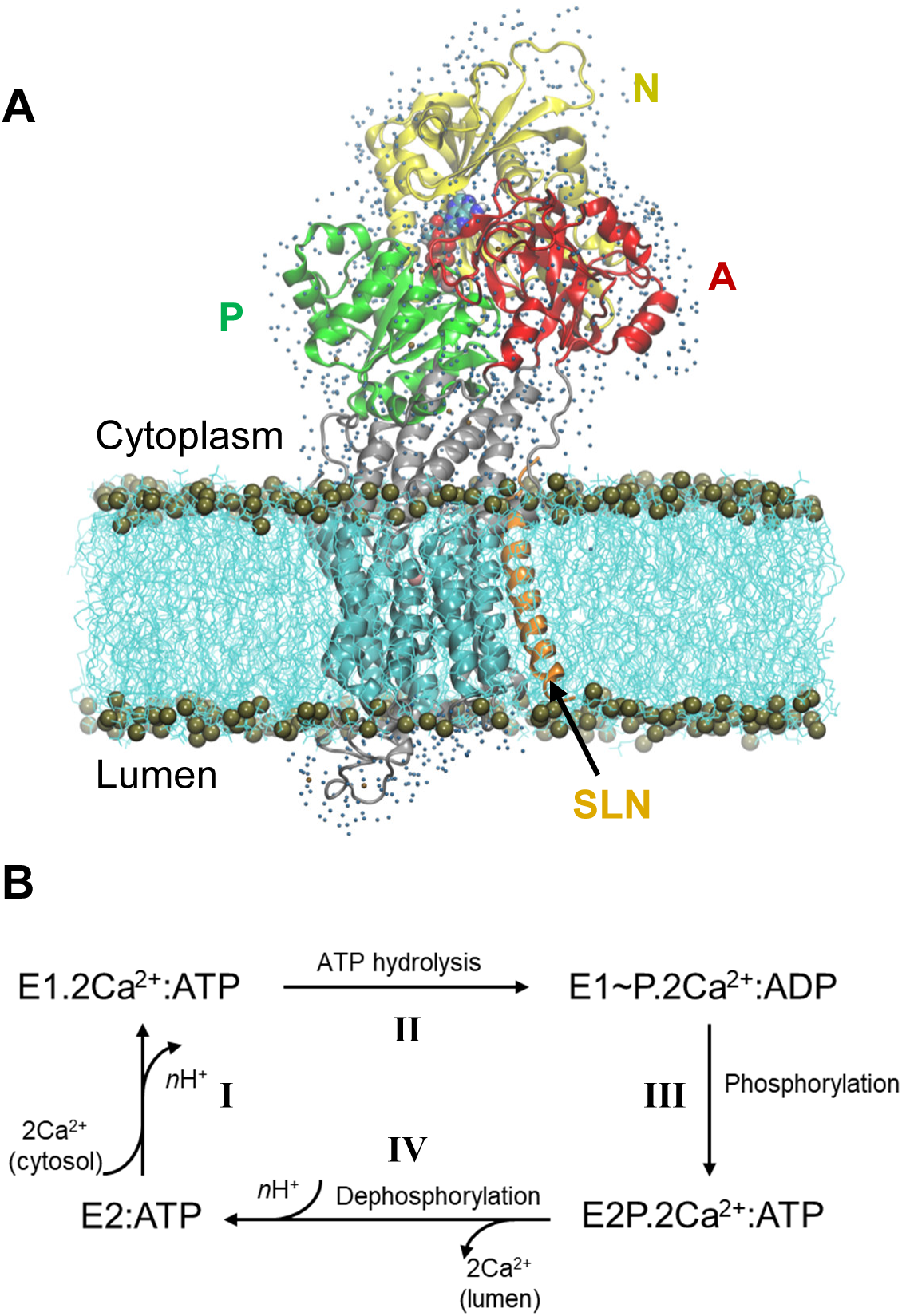
Structure of SERCA1a in the presence of sarcolipin (SLN) and the cycle of Ca^2+^ transport. (A) Structure of SERCA1a-SLN embedded in a POPC membrane, with one shell of water surrounding its cytosolic and luminal parts. The N-, P- and A-domains are in yellow, green and red, respectively, the TM domain in cyan and the rest of the protein in gray. SLN is in orange and ATP is in spheres colored according to atom name, C: cyan, N: blue, O: red, P: tan and Mg^2+^: pink. The membrane is in cyan lines with phosphorus atoms in tan spheres. Finally, water molecules are blue dots. (B) Simplified cycle of the Ca^2+^ transport from the cytoplasm to the sarcoplasmic reticulum lumen.

During Ca^2+^ transport across the sarcoplasmic reticulum membrane, SERCA1a undergoes conformational changes from the Ca^2+^ high-affinity state (E1) to the Ca^2+^ low-affinity state (E2). The functional cycle may be described in four main steps (Figure 1B, adapted from Møller *et al*. (5)): **I-** SERCA1a in the presence of ATP binds two cytoplasmic Ca^2+^ ions to form the **E1.2Ca^2+^:ATP** complex, in which the two Ca^2+^ binding sites are occluded (PDB codes: 1vfp (6), 3ar2 (7), 1t5s (8) and 3tlm (9), see Table S1 in Supplemental Information for details about the structures). At this step, two or three protons, H^+^, are released from the calcium binding sites into the cytoplasm. **II-** Following the occlusion of the Ca^2+^ binding sites, ATP is hydrolyzed to ADP, leading to the formation of a highly energetic intermediate, **E1∼P.2Ca^2+^:ADP** (PDB codes: 2zbd (10) and 1t5t (8)). **III-** The protein is then autophosphorylated and switches to the E2 conformation. The ADP molecule is exchanged with ATP and the Ca^2+^ binding sites open toward the lumen, resulting in the **E2P.2Ca^2+^:ATP** complex (PDB codes: 5a3r (11) and 3b9b (12)). **IV-** The two Ca^2+^ ions are then released into the lumen, accompanied by the protonation of *n* acidic residues (2 ≤ *<u>n</u>* ≤ 3) of the Ca^2+^-binding sites. In this step, auto-dephosphorylation produces the **E2:ATP** complex (PDB code: 3w5c (13)), where a structural rearrangement of the transmembrane helices slightly opens the protonated Ca^2+^-binding sites toward the cytoplasm, inducing protons release and enabling the protein to bind cytoplasmic Ca^2+^ again, as in step I. Since an ATP analog is always present in the structures that we consider in this study (except for 3w5c that will be discussed below), from here on ATP will be omitted from the notation of the states.

The existence of an E1 free-of-Ca^2+^ intermediate state was hypothesized between E2:ATP and E1.2Ca^2+^:ATP states, with the Ca^2+^-binding sites open toward the cytoplasm (13–15). In this hypothesis, it is unclear if the Ca^2+^-binding sites are free of any ions or if Mg^2+^ or K^+^ ions may replace Ca^2+^. The structure of this intermediate is still unknown, but those of SERCA1a in a putative intermediate E1 state have been proposed in complex with regulatory peptides, like sarcolipin (SLN) (PDB codes: 3w5a (13) and 4h1w (15)) or phospholamban (PLN) (PDB code: 4kyt (12)). In these complexes, SERCA1a is in an E1-like state with a large opening of the gate between the calcium binding sites and the cytoplasm.

The regulatory transmembrane peptide, SLN, composed of 31 residues, is mostly expressed in skeletal muscle cells (16, 17). Although it is widely accepted that SLN is an inhibitor of SERCA1a, the detailed mechanism of this inhibition remains unknown. The numerous experimental studies of the effect of SLN on SERCA1a clearly show a moderate decrease in Ca^2+^ affinity, whereas contradictory results are obtained concerning the ATP hydrolysis rate, most probably due to the differences in the studied biological systems ((18) and references herein).

The two crystal structures of the SERCA1a-SLN complex available in the Protein DataBank (PDB codes: 3w5a (13) and 4h1w (15)) were resolved in the presence of Mg^2+^. One Mg^2+^ is bound in the Ca^2+^ binding sites in 3w5a and two Mg^2+^ in 4h1w. Nonetheless, SLN binds to SERCA1a in a similar way, in a groove formed by transmembrane helices TM2, TM6 and TM9 (Figure S1 in Supplemental Information). The two structures are very similar in their transmembrane part, with a root-mean-square deviation (RMSD) over the TM C_α_ atoms of 0.6 Å, whereas the cytoplasmic domains are more open in 3w5a than in 4h1w, yielding to an RMSD over all C_α_ atoms of 2.2 Å.

As described by Toyoshima *et al.* (13) and Winther *et al.* (15), the most remarkable difference between the crystal structures of SERCA1a (E1.2Ca^2+^, 1vfp), and SERCA1a-SLN (SERCA1a.Mg^2+^:SLN, 3w5a, and SERCA1a.2Mg^2+^:SLN, 4h1w) is the opening of a large mouth in SERCA1a-SLN that leads to the Ca^2+^ binding sites. The opening of this mouth is questioning since, by making these sites more accessible, it is expected to impact the ATPase activity by increasing the Ca^2+^ uptake affinity, while experimental observations show a decrease in the apparent Ca^2+^ affinity. However, this large mouth does not seem to be only due to the presence of SLN, since it is also observed in the structure of SERCA1a.Mg^2+^ in the absence of SLN (3w5b).

In addition to X-ray diffraction experiments, *in silico* experiments were performed to investigate at a molecular level the role of SLN on SERCA1a. This role has been investigated by all-atom MD simulations (19–21), where analyses were focused on the Ca^2+^ binding sites and on the cytosolic part of TM4 (helix M4S4 spanning residues P312-K329). From these analyses, it was concluded that the presence of SLN perturbs the Ca^2+^ binding sites occlusion at E309, leading to an increase in the distance E771-D800, which produces incompetent Ca^2+^ binding sites and allows Ca^2+^ backflux to the cytosol.

Here, we use another *in silico* method, namely Normal Modes Analysis (NMA), to decipher the mechanism of inhibition of SERCA1a by SLN and to answer the question if this inhibition is due to Ca^2+^ binding, to autophosphorylation or to both. In the NMA method, the dynamics of a protein is approximated by the sum of its internal vibrations. Each normal mode (NM) is a vector corresponding to the direction of the vibration, and the lowest-frequency modes correspond to large-amplitude motions, which are usually equivalent to the concerted motions of the structure. The main interest of NMA is to provide a direct access to the principal component motions (the most collective movements), so it is a method of choice to study large conformational changes of proteins. To investigate the effects of Sarcolipin on the SERCA1a structure and motion, we chose to study and compare two systems starting from the same PDB structure, SERCA1a.Mg^2+^:SLN (3w5a): one in the presence of SLN, E1.Mg^2+^:SLN, and one from which SLN was removed, E1.Mg^2+^ (see Methods for details). The choice of starting from the same PDB structure, and not from the two crystal structures with and without SLN (3w5a and 3w5b, respectively), was made to enable us to identify the only effects of SLN, without any other considerations, like the difference of biological materials from which SERCA1a was obtained. Indeed, while SERCA1a was extracted from rabbit muscle for the resolution of the SERCA1a.Mg^2+^:SLN (3w5a) structure, it was expressed as a recombinant rabbit protein in *Chlorocebus Sabaeus* (COS7) cells for the SERCA1a.Mg^2+^ (3w5b) structure (13). Contrary to previous studies on the SERCA1a state-transition, where coarse-grained NMA methods were used − with either only C_α_ atoms (22) or rigid-blocks residues (23) −, and where the effect of the membrane was not taken into account, here we consider the all-atom model. The protein is embedded in a 1-palmitoyl-2-oleyl-sn-glycero-3-phosphocholine (POPC) bilayer membrane, and its solvent accessible parts are surrounded by one water shell. To our knowledge, this is the first NMA study of a membrane protein embedded in an explicit membrane, where the entire system is considered as an all-atom model.

## Methods

### Preparation of the crystal structure

#### Crystal structures

The two SERCA1a-SLN structures available in the Protein DataBank (https://www.rcsb.org/), code: 3w5a (13) and 4h1w (15), were resolved in the presence of millimolar concentrations of Mg^2+^ instead of Ca^2+^ to stabilize a Ca^2+^-deprived state of SERCA1a (40 mM of MgSO_4_ for the former and 75 mM of MgSO_4_ for the latter). They are similar, however, the 3w5a structure has been obtained with a slightly better resolution than 4h1w (3.01 Å and 3.10 Å respectively). In addition, in the 4h1w structure, there are some missing residues in the SERCA1a luminal part, i.e. in the L1-2 loop (residues 79 to 86) and in the L8-9 loop (residues 883 to 887). Therefore, we chose to use 3w5a coordinates for this study. In this structure, two chains are available for SERCA1a (chains A and B), but SLN (chain C) is clearly embedded in chain B. Therefore, only chain B (SERCA1a) and chain C (SLN) were considered.

#### Positioning of the adenosine triphosphate (ATP)

In the 3w5a crystal structure, in the ATP site, a trinitrophenyl adenosine monophosphate (TNPAMP) is bound, and a Mg^2+^ ion, chelated by only 4 water molecules, is present in its vicinity. To replace TNPAMP with ATP, the crystal structure of SERCA1a in its E1.2Ca^2+^ state was used (PDB code: 1vfp (6)). In this structure, an adenosine-[β,γ-methylene]triphosphate (AMPPCP), an ATP analog, is bound in the ATP pocket. A Mg^2+^ ion is chelated by AMPPCP and 2 water molecules. To do the replacement, in 3w5a, all residues with at least one heavy atom within 5 Å from TNPAMP-Mg^2+^ were selected, and the structure 1vfp was superimposed on these residues. We observed that by this superimposition AMPPCP-Mg^2+^ occupied the same location as TNPAMP-Mg^2+^. Therefore, these coordinates, in addition to those of the two Mg^2+^-chelating waters, were copied to replace those of TNPAMP-Mg^2+^ with their four Mg^2+^-chelating waters. Then AMPPCP was changed into ATP by replacing the β,γ-methylene with an oxygen atom. To adjust the bond lengths between the oxygen atom and the phosphate groups, the energy of the β,γ-phosphates was minimized by 10 steps of steepest descent method while the rest of the system was kept fixed.

#### Additional ions and crystal water molecules

In the 3w5a structure, an additional Mg^2+^ ion, chelated by two water molecules, sits in the calcium binding sites. A Na^+^ ion is chelated by L711, K712, A714 and E732. This monovalent cation is present in almost all SERCA1a structures and plays a stabilizing role in the P-domain Rossman fold (5). These additional ions (Mg^2+^ and Na^+^) and the 2 crystal water molecules that chelate the Mg^2+^ ion were also kept in this study.

#### Cleaning the structure

Since in a crystal structure at a resolution R > 1 Å, the C, N and O atoms are not distinguishable from each other, all asparagine, glutamine, and histidine residues were carefully examined to define their orientation. In addition, the pKa of the charged residues were calculated to determine their protonation state. All this was done using the protein preparation wizard from the Schrödinger Suite (https://www.schrodinger.com/).

#### Membrane and one layer of water

Based on this prepared structure, two systems were considered, one in the presence of SLN, and one where SLN was removed. For both these systems, the transmembrane part of the protein was embedded in a 1-palmitoyl-2-oleyl-sn-glycero-3-phosphocholine (POPC) bilayer using the Membrane Builder module (24–26) from the CHARMM-GUI webserver (27) and the Orientation of Proteins in Membrane (OPM) database (28). POPC lipids were chosen because: first, phosphatidylcholine (PC) headgroups are the major phospholipid component (70% of the phospholipid content) of the SR membrane and second, palmitoyl and oleoyl acyl chains represent 40% of the fatty acyl chains content of the SR (29, 30). Moreover, POPC lipids assemble in a bilayer with a thickness of about 40 Å in agreement with a functionally optimal membrane thickness for SERCA1a (31). The bilayer membrane was made of 376 lipids for SERCA1a-SLN and 379 lipids for SERCA1a alone. The solvation and ionization of the systems were also made using the CHARMM-GUI server. For this, in each system, the two soluble parts of the protein, on the luminal and cytosolic sides, were surrounded by a water box with randomly distributed K^+^ and Cl^-^ ions to reach a concentration of 100 mM. Then, for energy minimizations, only one layer of water and ions around the protein (within 2.8 Å from heavy atoms) was conserved to prevent the artefact shrinkage of the protein due to energy minimizations. Regarding this solvation shell, two systems were considered for the complex SERCA1a-SLN: one in which this water layer was created randomly (it consists of 1346 water molecules, 9 K^+^ and 2 Cl^-^), and one in which this layer was copied as it from the system of the isolated SERCA1a (1346 water molecules, 6 K^+^ and 4 Cl^-^). Despite the difference of charges the results were similar. Therefore, in the article only the results with the same water shell around SERCA1a-SLN and SERCA1a (1346 water molecules, 6 K^+^ and 4 Cl^-^) are presented. In this case, except for the 3 extra lipids that compensate the absence of SLN in SERCA1a, the only difference between the two systems is the presence or absence of Sarcolipin. All hydrogen atoms were created by the CHARMM-GUI website.

### Calculations

#### Potential energy force field

All the following calculations were carried out with program CHARMM (32) using the CHARMM36 all-atom force field (33, 34), excluding the CMAP correction, which is not needed for energy minimizations and normal mode calculations. The TIP3P model (35, 36) was used for water molecules. Electrostatic and van der Waals interactions were switched to zero with a cut-on distance of 6 Å and a cutoff distance of 10 Å and the relative dielectric constant was equal to 3. These values were chosen to avoid important distortions of the structure due to extreme energy minimizations.

#### Energy minimizations and normal modes calculations

Each system was energy minimized by 1000 steps using the steepest descent method followed by conjugate gradient method for thousands of additional steps until the root mean square of the energy gradient (GRMS) fell below 10^-4^ kcal/mol.Å^-1^.

The two energy-minimized structures were named according to their state, E1.Mg^2+^:SLN and E1.Mg^2+^, to avoid confusion with the crystal structures in presence of Mg^2+^, namely SERCA1a.Mg^2+^ (3w5b), SERCA1a.Mg^2+^:SLN (3w5a) and SERCA1a.2Mg^2+^:SLN (4h1w). E1.Mg^2+^:SLN and E1.Mg^2+^ were considered as starting systems to calculate normal modes. Because of the large number of atoms in each system (> 70,000 atoms) the DIMB method (Diagonalization in a Mixed Basis) (37, 38) was used in program CHARMM. But to be able to perform the calculations in a reasonable time (< 10 days), we accelerated this method by making it work on a GPU system, in a home-made adaptation of CHARMM. This adaptation was tested beforehand on small proteins to insure that the results were the same as those obtained on CPU. For each system 206 modes were calculated, the 6 trivial global translation-rotation modes with null frequencies, and 200 non-trivial modes (modes 7 to 206), sorted according to the ascending order of their frequencies. The modes were considered as converged when the maximum eigenvectors convergence was less than 0.03.

##### Modes overlaps

The percentage of overlap (*p*) between two vectors, 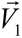 and 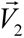 was calculated using the following equation:

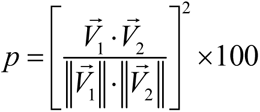

where 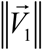 and 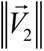 are the norms of vectors 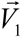 and 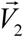.

In this study we calculated the percentage of overlap (or projection) between the modes of a structure that we call *A* and the difference coordinates vector between structures *A* and *B*. So 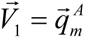, where 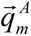 is mode *m* of structure *A*, with *m*∈[7,206], and 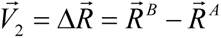, where 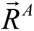 and 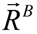 are the coordinates of structures *A* and *B*, respectively. Vector 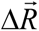 may also be written *A* → *B*. Prior to the projection, structure *B* is superimposed on structure *A* using C_α_ atoms. The projection is also done considering only C_α_ atoms in both vectors,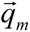 and 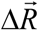, in order to avoid misleading directions due to sidechains. The higher the percentage of overlap between these two vectors (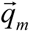 and 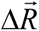), the most straightforward is the transition from *A* to *B*. Here *A* refers to one of the two energy-minimized structures (E1.Mg^2+^ and E1.Mg^2+^:SLN) and *B* to one of the two chosen crystal structures (1vfp for state E1.2Ca^2+^ and 3w5c for state E2). For the E1.2Ca^2+^ state, 1vfp was preferred to the other structures, 3ar2, 1t5s and 3tlm, despite its slightly lower resolution in some cases, because 1t5s and 3tlm lack the disulfide bridge that stabilizes the protein, and the ATP analog in 3ar2 chelates Ca^2+^ instead of Mg^2+^ (see Table S1 in Supplemental Information). For the E2 state, the structure of 3w5c, although lacking an ATP analog, was preferred to 2dqs, because it is free of exogenous molecules like thapsigargin. 2dqs, which has the ATP analog, also contains thapsigargin to stabilize it by disrupting the communication between the transmembrane domain and the cytosolic headpiece of the ATPase (39, 40). However, despite these differences, the two E2 structures, 3w5c and 2dqs, are very similar, with an RMSD between all their C_α_ atoms of only 0.6 Å.

We also calculated the percentage of overlaps between all the modes calculated from one structure (*B*) over one chosen mode calculated from another structure (*A*), in which case,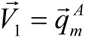 and 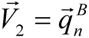, where *m* = 54 when *A* = E1.Mg^2+^:SLN and *n* ∈[7,206] for *B* = E1.Mg^2+^, or *m* = 40 when *A* = E1.Mg^2+^ and for *B* = E1.Mg^2+^:SLN (see subsection 2 in the *n* ∈[7,206] Results).

##### Atomic fluctuations

For each structure, E1.Mg^2+^:SLN and E1.Mg^2+^, the atomic fluctuations, *f_i_*, were calculated from the 200 modes using:

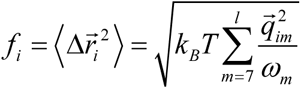

where 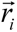 is the position of atom *i*, 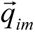 its component in mode *m*, ω*_m_* the frequency of mode *m*, *T* is the temperature and *k_B_* the Boltzmann constant. The sum runs from 7 to *l*, which is usually equal to 3*N*, where *N* is the number of atoms. However, the fluctuations are dominated by those of the lowest frequency modes, since *f_i_*^2^ is inversely proportional to the frequency. Here *l* = 206. Because even when all modes are considered, the fluctuations obtained from NM in the all-atom model are undervalued compared to those obtained from molecular dynamics or the crystal B-factors, for this calculation *T* was taken equal to 1300 K.

The fluctuations were also calculated from the crystal structure B-factors using the equation:

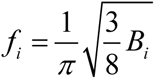

##### Correlations

The correlation *C_ij_* between two C_α_ atoms, *i* and *j*, calculated from one mode, *m*:

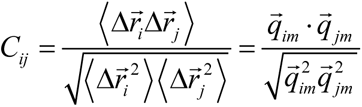

##### Analyses and visualization

All the analyses were performed with CHARMM. The plots were drawn using Kaleidagraph (http://www.synergy.com) and the images were done using VMD (Visual Molecular Dynamics, https://www.ks.uiuc.edu/Research/vmd/ (41)).

## Results

Since it has been proposed that the crystal structure of the SERCA1a-SLN complex is in an intermediate state between the Ca^2+^-deprived state (E2) and the Ca^2+^-bound state (E1.2Ca^2+^), the role of SLN in the conformational transition of SERCA1a toward E2 or E1.2Ca^2+^ was first investigated.

### 1-E1.Mg2+:SLN has the propensity to go toward E2 and E1.Mg^2+^ toward E1.2Ca^2+^

In all this study, the reference structure for the E1.2Ca^2+^ state was chosen to be 1vfp (chain A) and for the E2 state, 3w5c. The 200 lowest-frequency modes of each energy-minimized system, E1.Mg^2+^:SLN and E1.Mg^2+^, were projected on the difference coordinates vector with either E1.2Ca^2+^ (1vfp) or E2 (3w5c). The results are presented in Figure 2, where the percentages of overlaps are plotted *vs* the NM frequencies. We observe that the lowest-frequency modes, which usually correspond to the internal most collective motions of a protein, do not present high overlaps with the difference vectors. This is because these modes are only collective when considering the entire system, i.e., the protein plus the membrane and the water shell. Here these modes mainly correspond to the global vibrations of the rectangular membrane, in which the transmembrane domain is embedded. Therefore, the TM domain follows these vibrations, in addition to a global motion of the three cytosolic domains with respect to the membrane. When we consider the modes of slightly higher frequency, global internal motions of the protein itself are observed, resulting in higher overlaps.

**Figure 2.**
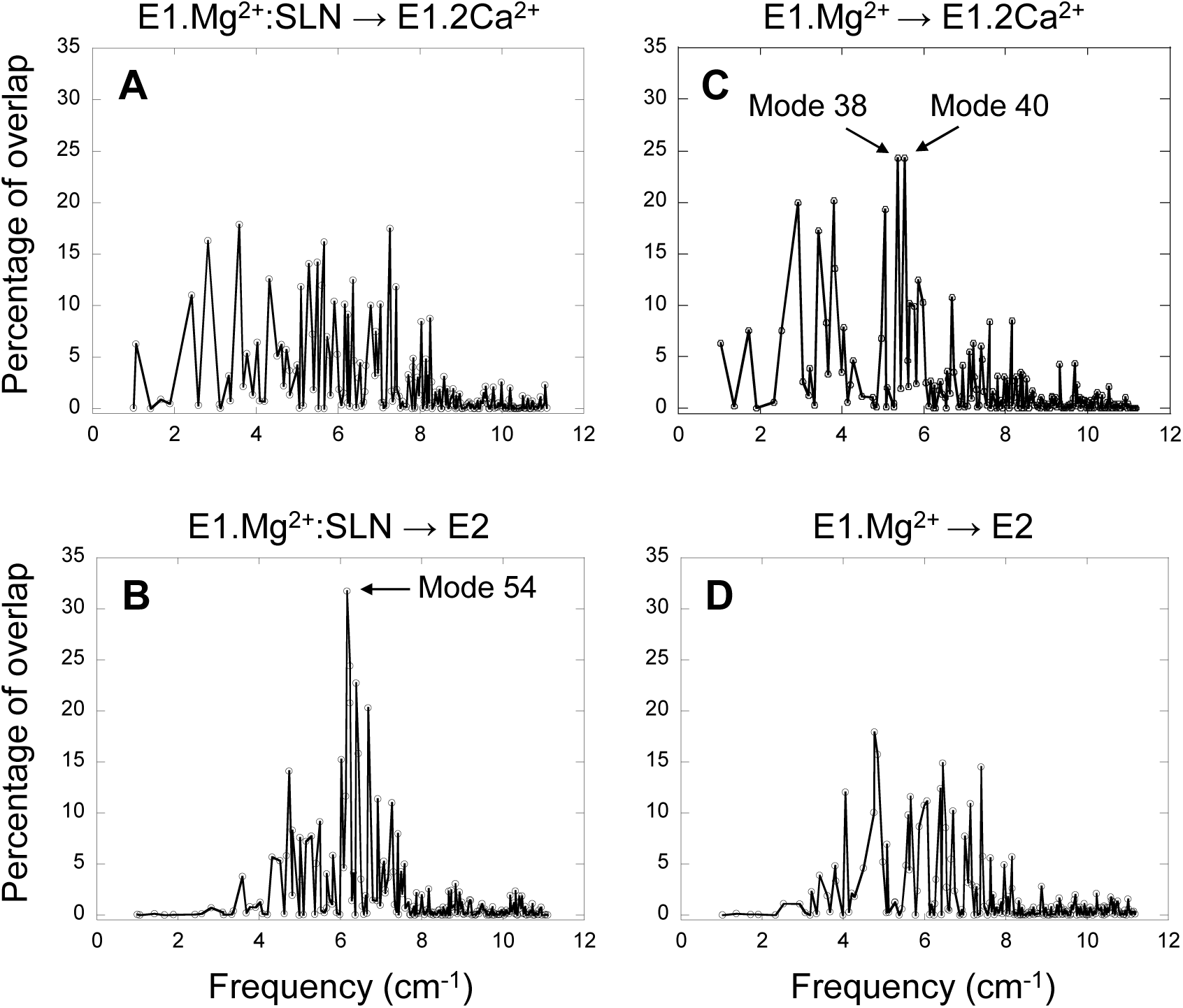
Projection of the normal modes on the difference coordinates vectors. Percentage of overlaps between the 200 NMs of E1.Mg^2+^:SLN and the difference vectors E1.Mg^2+^:SLN → E1.2Ca^2+^ (A) and E1.Mg^2+^:SLN → E2 (B). Percentage of overlaps between the 200 NMs of E1.Mg^2+^ and the difference vectors E1.Mg^2+^ → E1.2Ca^2+^ (C) and E1.Mg^2+^ → E2 (D). E1.Mg^2+^:SLN and E1.Mg^2+^ are energy-minimized structures and E1.2Ca^2+^ and E2 are crystal structures (PDB codes: 1vfp and 3w5c, respectively). Only C_α_ atoms are considered for these projections.

For the E1.Mg^2+^:SLN structure, when its modes are projected on vector E1.Mg^2+^:SLN → E2 the highest overlap is 32 %, obtained for mode 54, the frequency of which is 6.16 cm^-1^, whereas when projected on vector E1.Mg^2+^:SLN → E1.2Ca^2+^, the highest overlap is only 18 % (Figure 2A, B). This shows the preference of E1.Mg^2+^:SLN to go toward E2 than toward E1.2Ca^2+^.

Conversely, in the absence of SLN, for E1.Mg^2+^, the highest percentage of overlap, 24 %, is observed for two modes corresponding to motions toward E1.2Ca^2+^, modes number 38 and 40, with frequencies 5.38 cm^-1^ and 5.54 cm^-1^, respectively (the modes are ranked according to the ascending order of their frequencies), whereas, toward E2, the highest overlap was only 18 % (Figure 2C, D). This shows a higher tendency of E1.Mg^2+^ to go toward E1.2Ca^2+^ than toward E2.

To decipher the role of the different domains of SERCA1a in the conformational changes required for the transition toward the E1.2Ca^2+^ or E2 states, the three modes that gave the highest percentage of overlap (mode 54 of E1.Mg^2+^:SLN and modes 38 and 40 of E1.Mg^2+^) were further analyzed.

### 2-The P-domain plays a central role in the transition toward the E2 state

The motion of E1.Mg^2+^:SLN along mode 54, which brings the structure toward E2, is shown in Figure 3A. It corresponds to the rotation of the A-domain around the region of the P-domain which is in contact with it. The P-domain rotates in the opposite direction around the TM5 helix, and the N-domain, which is in the continuity of the P-domain, rotates in the same direction. The TM domain does not present any wide motions, except for the N-terminus of helix TM2, which is in contact with the C-terminus of SLN and moves in concert with it.

**Figure 3.**
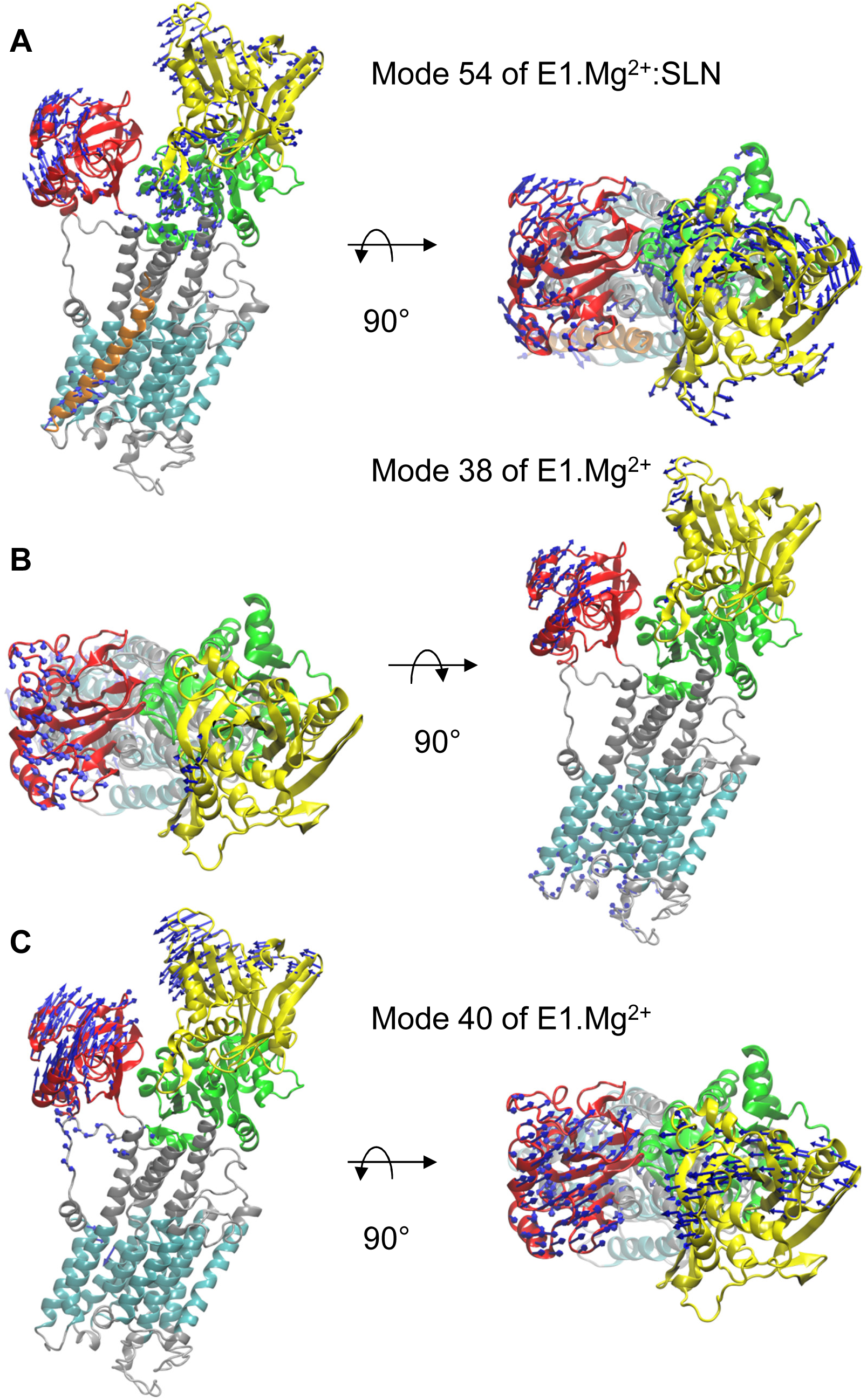
Motions along the three modes identified from the overlaps. In each of the three panels a profile view and a top view of the protein are presented. The same color code as in Figure 1A is used. The directions of motions are shown as blue arrows. The lengths of the arrows are proportional to the amplitude of displacement of the C_α_ atoms when the entire system is displaced by mass-weighted RMSD of 3 Å. For clarity, the arrows corresponding to displacement amplitudes under 3 Å are omitted. The motions are shown along mode 54 of E1.Mg^2+^:SLN (A) and along modes 38 (B) and 40 (C) of E1.Mg^2+^.

To further describe the concerted motions of the different domains of SERCA1a for mode 54, the correlations from mode 54 of E1.Mg^2+^:SLN were calculated (Figure 4A). The heatmap of these correlations shows two compact regions, the TM5 to TM10 sub-domain and the P-domain, that move in a concerted way (although with different amplitudes as observed in Figure 3A). Whereas the TM5-TM10 sub-domain is mainly correlated with the P-domain, the P-domain is also correlated with the N-domain and anticorrelated (concerted in the opposite direction) with TM1, TM2 and the A-domain. TM1 and TM2 are two of the three helices that link the A-domain to the rest of the protein.

**Figure 4.**
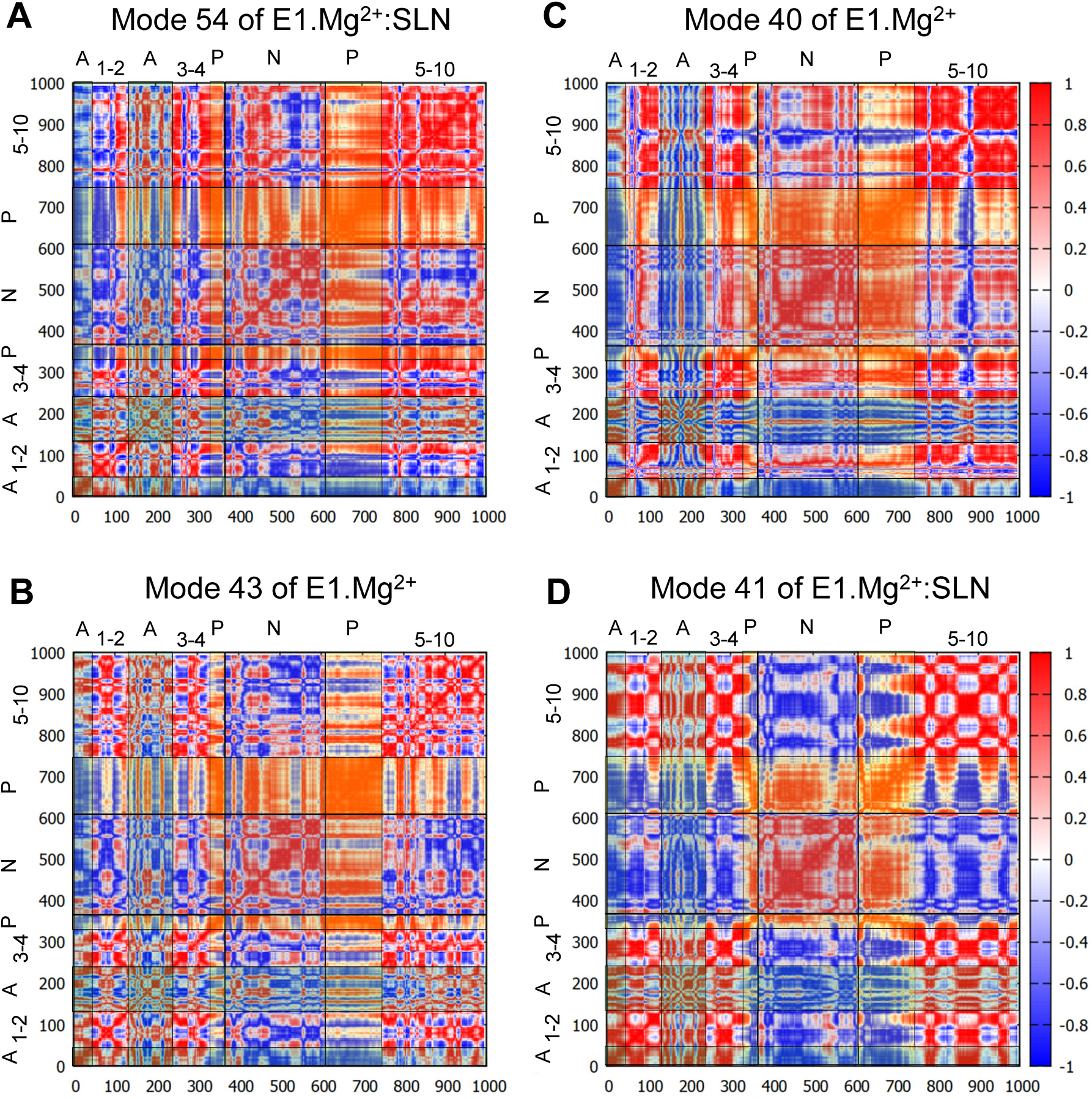
Correlation heatmaps. The correlations are calculated from modes 54 (A) and 41 (D) of E1.Mg^2+^:SLN and modes 40 (C) and 43 (B) of E1.Mg^2+^. The red color corresponds to high correlations, white to the absence of correlations and blue to high anticorrelations. However, to facilitate the delimitation of the cytosolic domains, transparent vertical and horizontal strips were overlaid on each heatmap, with the following color code: N-domain: gray, P-domain: yellow and A-domain cyan, which modifies the perception of the correlations colors. In addition the names of the domains are added on the top and the left of each heatmap, where the numbers refer to the TM helices.

Does a similar motion exist in the absence of SLN, i.e., in E1.Mg^2+^, that was not detected with the projections on the difference coordinates vector? To answer this question, we projected all the calculated modes of E1.Mg^2+^ on mode 54 of E1.Mg^2+^:SLN. The highest percentage of overlap is 60 % for mode 43 of E1.Mg^2+^ (frequency = 5.65 cm^-1^). However, this mode presents only a percentage of overlap of 12 % with the difference vector E1.Mg^2+^ → E2. The analysis of this mode shows similar motions of the protein as those along mode 54 of E1.Mg^2+^:SLN, but with a smaller amplitude and involving less extended regions of the protein (Figure S2A in Supplemental Information). The heatmap of the correlations calculated from mode 43 of E1.Mg^2+^ (Figure 4B) shows that the P-domain is still the only one that presents global correlations with the rest of the protein. However, the correlation coefficients are smaller than those calculated from mode 54 of E1.Mg^2+^:SLN, due to slight differences in the directions of motion.

In conclusion, in the transition toward E2, first, the P-domain is the only compact region that presents global correlations, whether they be positive or negative, with the rest of the protein; and second, while in the presence of SLN, these correlations are strong, coinciding with a good propensity for going toward E2, in the absence of SLN these correlations are decreased as well as the propensity for the E2 transition. So we conclude that the P-domain plays a central role in the transition toward E2.

### 3-A close correlation between the P- and N-domains is important for the transition toward the E1.2Ca^2+^ state

In the absence of SLN, i.e., for structure E1.Mg^2+^, two modes have been identified for the transition toward E1.2Ca^2+^, modes 38 and 40 (Figure 2C). The motion along mode 38 is mainly located in loops (Figure 3B), and therefore cannot drive the state transition. So, despite its overlap, this mode will not be considered in the rest of the manuscript. The motion along mode 40 is collective and resembles a hinge-bending motion that brings the A- and N-domains close to each other, like in the E1.2Ca^2+^ state. In addition, it brings the kinked helix TM1 − and more specifically residue L60, the junction between the two parts of TM1 − near TM4, starting to close the protein mouth. These data highlight the concerted motion between the cytoplasmic domains and the occlusion of the Ca^2+^ binding sites, as previously suggested from structural analyses (6, 8).

The heatmap of the correlations calculated from mode 40 (Figure 4C) shows that in this case, the P- and N-domains act together as a concerted subunit, which is anticorrelated with the A-domain. This result is consistent with the hinge-bending-like motion observed for this mode (Figure 3C).

To investigate whether a mode of E1.Mg^2+^:SLN with a motion similar to that of mode 40 of E1.Mg^2+^ exists, all modes of E1.Mg^2+^:SLN were projected on mode 40 of E1.Mg^2+^. The best percentage of overlap (48 %) was obtained for mode 41 (frequency 5.50 cm^-1^) of E1.Mg^2+^:SLN. Although this mode has almost the same frequency as mode 40 of E1.Mg^2+^, the motions of the cytoplasmic domains along it are of smaller amplitudes and they are more localized (Figure S2B in Supplemental Information). Its moderately low frequency is probably due to a relatively large amplitude motion of the TM domain along the normal to the membrane plane. This motion of the TM domain is not observed in the transition toward the E1.2Ca^2+^ state. The correlation heatmap corroborates this observation, where the P- and N-domains are less correlated with each other, but highly anticorrelated with all the rest of the protein (Figure 4D). Therefore, the presence of SLN seems to hamper the transition toward E1.2Ca^2+^, because at the same time it decreases the correlation between the P- and N-domains and it correlates the closing hinge-bending motion with an upward movement of the TM domain, which is not necessary for this transition.

### 4-SLN reduces the flexibility of the Ca^2+^ gating residues

The fluctuations of the C_α_ atoms of SERCA1a were obtained from the 200 modes that were calculated for each system. These fluctuations show that, for both systems, the cytoplasmic A-, N- and P-domains of the protein fluctuate more than its transmembrane domain (Figure S3 in Supplemental Information), reflecting the degree of accessibility to solvent of each region and therefore, its freedom of movement. We observe that these results are different from the fluctuations obtained from the thermal B-factors of the crystal structure of SERCA1a.Mg^2+^:SLN (3w5a), where they are reversed; in the PDB structure the TM domain is more fluctuating than the cytoplasmic domains due to the crystal packing. Indeed, in the crystal unit cell, the cytoplasmic domains are in close contact with other proteins of the cell, which is not the case of the TM domain. Therefore, the experimental fluctuations (B-factors) profile does not reflect the flexibility of SERCA1a.Mg^2+^:SLN in a membrane.

The fluctuation profiles in the presence and absence of SLN are very similar, so to observe the difference between them the curve of E1.Mg^2+^ was subtracted from that of E1.Mg^2+^:SLN. The difference curve (Figure 5A) shows that the observable modifications (> |0.2| Å) are very localized. In TM2, around residue F92, the fluctuations increase in the presence of SLN, because this region is in close contact with the peptide and follows its motion. The fluctuations of residues around F776, located in the C-terminal part of TM5, near the lumen, also increase in the presence of SLN; concomitantly, those of residue P784 in loop TM5-TM6, in the lumen, decrease. These two modifications of the fluctuations result from a slightly lesser structuration (less backbone-backbone H-bonds) in the bottom of TM5 (for F776), and a better structuration of the loop (for P784) probably due to the modification of the interactions with the neighboring TM6, consequently to the presence of SLN. In all the other regions (residues V304 in TM4, Q759 in TM5 and L807-G808 in TM6) the fluctuations decrease in the presence of SLN. Q759, L807 and G808 are in close contact with each other, and the reason of the decrease of their fluctuations will be given below in subsection 7. The most interesting residue is V304, because it is part of the Ca^2+^ binding site II, and represents, with E309, its opening gate. This needs a more thorough analysis.

**Figure 5.**
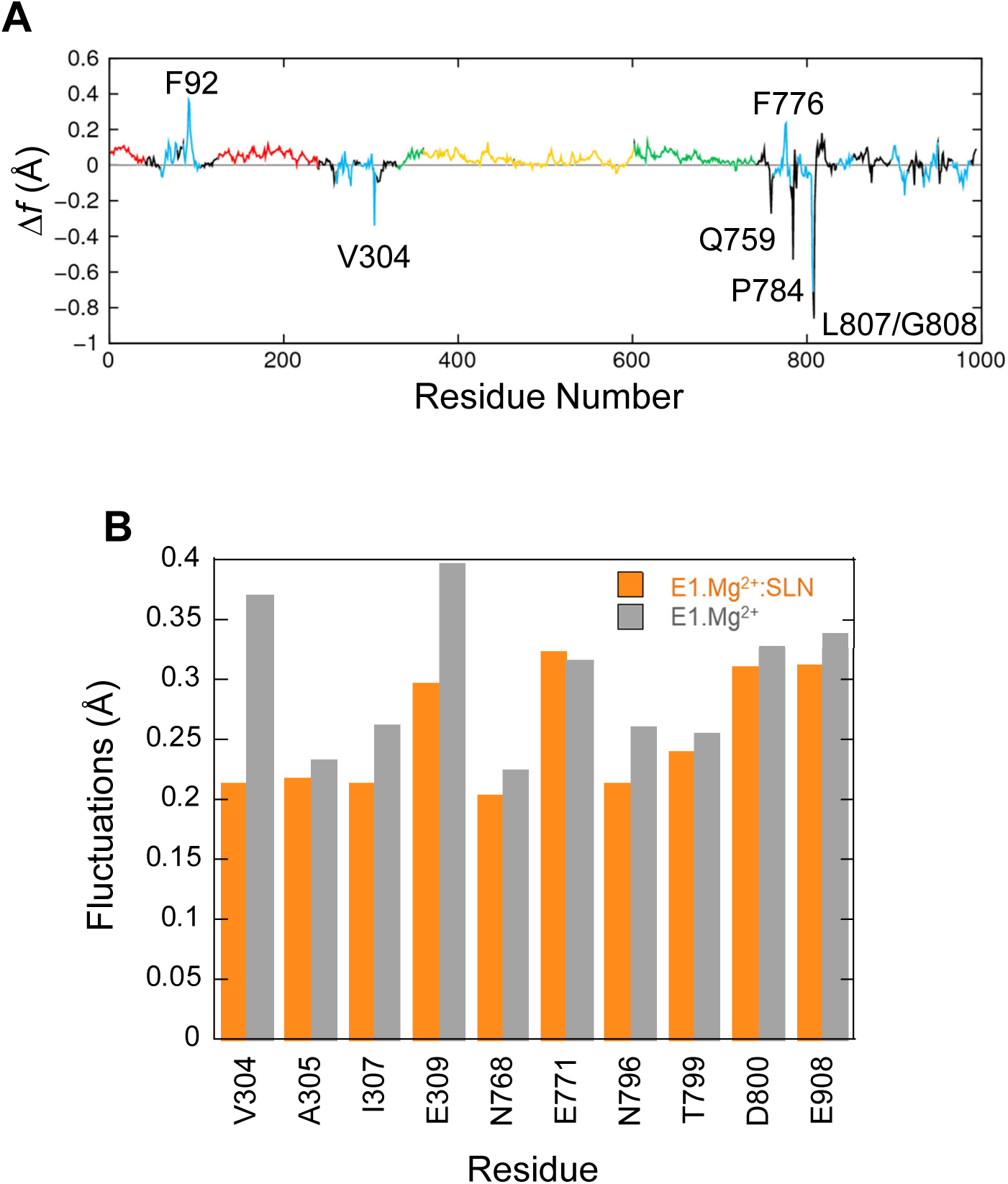
Fluctuations of residues. (A) Δ*f*, the difference of fluctuations of the C_α_ atoms calculated from the 200 modes. The fluctuations of E1.Mg^2+^ were subtracted from those of E1.Mg^2+^:SLN. The same color code as in Figure 1A is used to delimit the domains. (B) Average fluctuations of the Ca^2+^ chelating groups, i.e., the carboxyl, hydroxyl or carbonyl groups of the residues in the two Ca^2+^ binding sites. Orange bars for E1.Mg^2+^:SLN and gray bars for E1.Mg^2+^.

In its E1 state, SERCA1a can bind two Ca^2+^ ions in two adjacent sites, I and II, located between four transmembrane helices, TM4, TM5, TM6 and TM8, as observed in the structure of the E1.2Ca^2+^ state, 1vfp, (Figure 6A). Although Ca^2+^ ions enter from site II, they bind sequentially, first to site I then to site II, as shown by the studies using ^45^Ca^2+^ isotopes (42, 43), and the ion is more tightly chelated in site I due to its higher electronegativity (see the legend of Figure 6).

**Figure 6.**
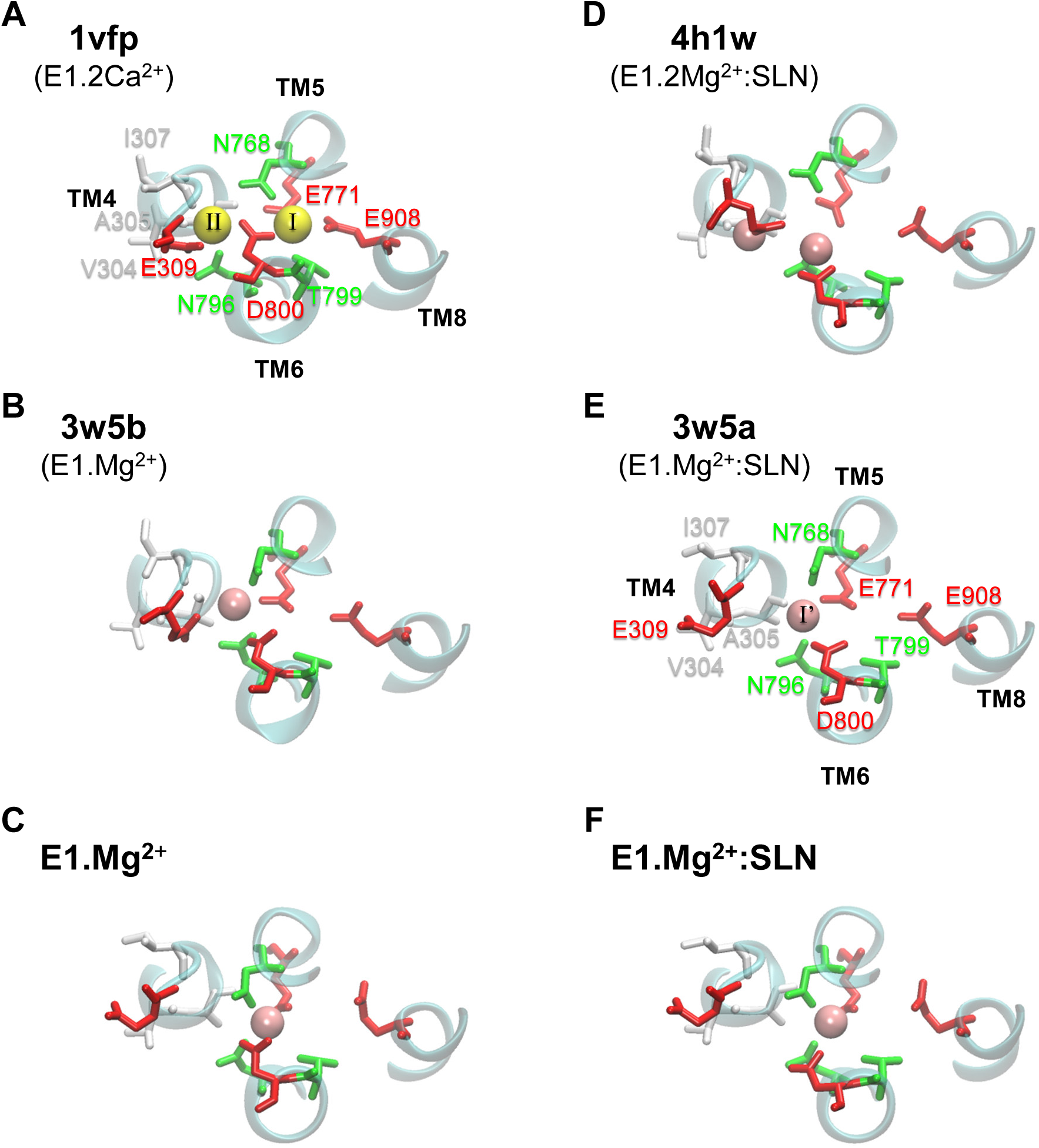
Ca^2+^ and Mg^2+^ binding sites in various SERCA1a structures. In (A, B, C) the structures are without SLN and in (D, E, F) they are in the presence of SLN. (A, B, D, E) show PDB structures, with the PDB code written in bold and the corresponding state given below the code in parentheses. (C, F) present the two energy-minimized structures, in the absence and presence of SLN, respectively. In each panel all the residues of the two Ca^2+^ binding sites are drawn, even when they are not chelating any ion, as for the structures with Mg^2+^. Ca^2+^ and Mg^2+^ ions are yellow and pink spheres, respectively. The binding sites residues are colored according to their type, acidic: red, polar: green and hydrophobic: white. (A) In site I, Ca^2+^ is chelated by eight oxygen atoms, of which four are from the carboxyl groups of acidic residues (TM5:E771, TM6:D800 and two from TM8:E908), two from the sidechains of polar residues (TM5:N768 and TM6:T799), in addition to two buried water molecules. In site II, Ca^2+^ is chelated by seven oxygen atoms, of which only three are from the carboxyl groups of acidic residues (two from TM4:E309 and one from TM6:D800), one is from the sidechain of a polar residue (TM6:N796) and three are from the carbonyl groups of the backbone of hydrophobic residues (TM4:V304, A305 and I307). (E) In 3w5a, Mg^2+^ sits in site I’, located between sites I and II and it is exclusively chelated by residues from these two sites, in addition to two water molecules. For clarity, water molecules are omitted and only small parts of helices TM4, TM5, TM6 and TM8 are drawn as cyan cartoons. Also for clarity, we chose to annotate only two structures, one with Ca^2+^ (1vfp) and one with Mg^2+^ (3w5a).

In the crystal structures with Mg^2+^ ions (3w5a, 3w5b, and 4h1w), the metal sits in an intermediate site, named I’, located between sites I and II, and encompassing a part of their residues (Figure 6B, D, and E). Site I’ is of a smaller volume than sites I and II, which makes it more appropriate to accommodate Mg^2+^ as mentioned by the authors of the two structures with SLN (13, 15).

Here we focus on the chemical groups of all residues in sites I and II that chelate Ca^2+^, because they include all residues of site I’ that chelate Mg^2+^. So we calculated for E1.Mg^2+^:SLN and E1.Mg^2+^ the average fluctuations of the carboxyl groups of E309, E771, D800 and E908, the hydroxyl group of T799, the sidechains carbonyl groups of N768 and N796 and the backbone carbonyl groups of V304, A305 and I307. The results are presented in Figure 5B. Only the fluctuations of the two gating residues, V304 and E309, clearly decrease in the presence of SLN. This difference of flexibility suggests that the presence of the peptide may hamper the access of the cation to its binding sites. This idea is supported by experimental observations where mutation of V304 by a more voluminous residue, V304L, slightly diminishes the Ca^2+^ binding affinity, although V304 only chelates Ca^2+^ by its backbone carbonyl (8). In addition, the identified mutations of the other gating residue, E309Q, E309D, E309A, E309K, E309L, E309M and E309F, generally prevent chelation of Ca^2+^ in site II and therefore significantly decrease the Ca^2+^ affinity (44–54). V304 is located at the C-terminus of the membrane part of helix TM4, and E309 in the hinge region (^308^PEG^310^) just before the N-terminus of the cytosolic part of TM4, also named helix M4S4. Due to the break of TM4, its cytosolic part (M4S4) is bent with respect to the plane of the membrane. In the presence of SLN, salt bridges are established between R324 and K328 at the C-terminus of M4S4 on the one hand, and SLN E2 on the other hand. Energy minimizations did not modify the bending angle of this helix, even in the absence of SLN. However, the salt bridges observed in the presence of SLN exerted a strain on M4S4. So V304 and E309, which are in the bending region on the opposite side of this helix, are probably hampered in their motion. In the absence of SLN, helix M4S4 is released, increasing the fluctuations of these two residues.

### 5-SLN weakens the chelation of Ca^2+^ at site I

Observation of the structures of the Ca^2+^ binding sites in the presence and absence of SLN after energy minimizations shows that they are similar except for residue D800 (Figure 6C, F). D800 is a key residue for calcium binding as, in the presence of Ca^2+^, it points toward both sites I and II to chelate the two calcium ions. In the crystal structure of SERCA1a.Mg^2+^:SLN (3w5a), which is the basis of this study, D800 is a little far for the chelation of Mg^2+^, the distance between the ion and the closest atom of D800 being 2.87 Å (Figure 6E). Toyoshima *et al.* (13) suggested that this could be the consequence of the approximate positioning of Mg^2+^ due to the low resolution (around 3 Å) of the crystal structure. But in fact D800 is far from site I, not only from Mg^2+^. This may be represented by the distance between the two attractive groups COO^-^ of D800 and NH_2_ of N768. N768 was chosen because it has the only sidechain with an attractive interaction with D800, while it is not located at the same helix (D800 is at TM6 and N768 at its facing helix TM5). More precisely, the distance was calculated between any oxygen atom, O_δ*_, of the D800 carboxylate group and atom N_δ2_ of the N768 amide group. This distance is 5.65 Å in the crystal structure of SERCA1a.Mg^2+^:SLN (3w5a), which is much bigger than that of E1.2Ca^2+^ (1vfp), where it is 3.08 Å (Table S2 in Supplemental Information). Besides, D800 does not point toward site II either. Nonetheless, in the presence of SLN, the relaxation of the structure by energy minimizations did not correct sufficiently this effect, since D800 did not point well toward site I, although the O_δ*_−N_δ2_ distance decreases to 4.83 Å, which is still more than 50 % larger than in the presence of Ca^2+^ (PDB structure 1vfp). Conversely, energy minimizations in the absence of SLN brought the O_δ*_−N_δ2_ distance to 3.40 Å, which is comparable to E1.2Ca^2+^ (3.08 Å).

In both E1.Mg^2+^:SLN and E1.Mg^2+^, Mg^2+^ is displaced close to E771 (distance 1.99 Å) due to energy minimizations. In the presence of SLN, D800 stays then far from the metal with a distance of 4.22 Å, whereas in the absence of SLN, it better adapts and comes to a distance of 3.92 Å, comparable with the distance in the crystal structure of SERCA1a.Mg^2+^, 3w5b, which is 3.69 Å. It therefore seems that it is more energetically demanding to D800 to come close to Mg^2+^ and to point toward site I in the presence of SLN than in its absence. The analysis of these energy-minimized structures shows that this may be due to the different structuration of helix TM6, to which belongs D800. Indeed, in the absence of SLN, TM6 kinks at 2/3 of its length, in the region ^800^DGLP^803^, because of the presence of a glycine and a proline, as reported for most of the structures of SERCA1a in the absence of SLN (5, 55, 56). The N-terminal part of TM6 (residue 789 to 800) is structured as an α−helix, whereas its C-terminal part (residues 801 to 810) comprises a 3-10 helix turn (Figure 7A). The α− and 3-10 helices are separated by an unstructured link, which lacks the intra-helix *i, i+*4 hydrogen bond between C=O of G801 and N-H of T805. This bond is replaced with a hydrogen bond between the carbonyl group of G801 and the hydroxyl group of the T805 sidechain. In the presence of SLN, the latter H-bond is broken and both C=O of G801 and OH of T805 sidechain establish hydrogen bonds with the facing N11 sidechain of SLN (Figure 7B). These new H-bonds result in a slight straightening of helix TM6 with a little rotation around its axis. Due to this constraint, residue D800, which neighbors G801 and T805, is not free anymore to point toward the Ca^2+^ binding site I. In the presence of Ca^2+^ instead of Mg^2+^, we assume that this constraint impedes D800 from adopting the right position for the chelation of one of the two calcium ions in the presence of SLN, resulting in a decrease of affinity for Ca^2+^.

**Figure 7.**
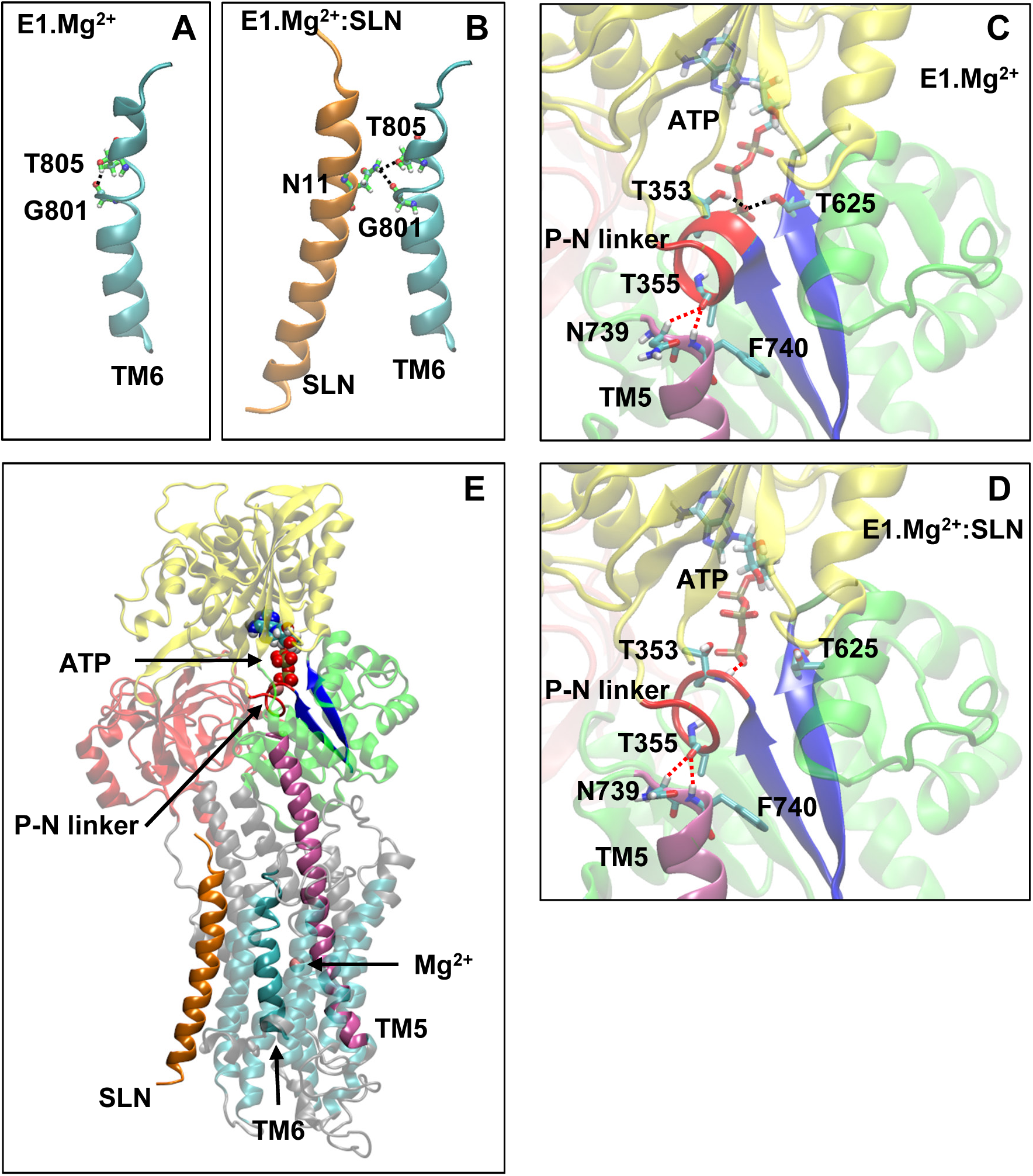
A few key structural modifications between E1.Mg^2+^ and E1.Mg^2+^:SLN. (A) and (B) Modification of the TM6 H-bonds in the absence and presence of SLN, respectively. (C) and (D) Modifications in the vicinity of ATP and the P-N linker in the absence and presence of SLN, respectively. D351 and Mg^2+^ of the ATP are not shown for clarity. The important H-bonds are in black dashed lines and some distances in red dashed lines. In both cases, the distances between the hydrogen and oxygen atoms that are related by these dashed lines are as follows: (C) in E1.Mg^2+^, T355-N739 = 3.3 Å, T355-N740 = 2.4 Å, ATP-T353 = 1.9 Å and ATP-T625 = 2.0 Å, and (D) in E1.Mg^2+^:SLN, T355-N739 = 2.5 Å, T355-N740 = 2.5 Å, ATP-T353 = 2.2 Å and ATP-T625 = 5.6 Å. (E) Zoom out where the protein is shown in transparency except for the elements that play an important role in the transmission of information from SLN to ATP to highlight their location. These elements are SLN, TM5, TM6, the P-N linker, the two β−strands that hold T625 and D351, ATP and the Mg^2+^ ion in the Ca^2+^ binding site. Generally, the same color code as in Figure 1A is used for the domains and for the explicit atoms. However, in (A) and (B) carbon atoms are in green to be distinguished from TM6, and in (C), (D) and (E), the P-N linker is in red, TM5 is in purple and the two β−strands that hold T625 and D351 are in dark blue.

### 6-The presence of SLN affects the phosphorylation site…

In addition to the Ca^2+^ binding sites, which are in its vicinity, SLN seems to also affect the phosphorylation site, which is distant by more than 35 Å. In the crystal structures, the ATP analog, when present, is positioned in the N-domain, with its phosphate groups pointing toward ^351^DKTG^354^, a motif common to all P-type ATPases, which comprises D351, the residue to be phosphorylated ((5) and references herein). The autophosphorylation of the protein consists of the formation of an aspartyl-phosphate through a nucleophilic association between the ATP γ-phosphate and the carboxyl group of D351 (57). To achieve the phosphorylation process, residues T353 and T625 (both in the P-domain) were described to interact through their side chains with the γ-phosphate to stabilize it. These two threonine residues are critical for proper orientation of the ATP γ-phosphate group preceding autophosphorylation (58–61). In the E1.2Ca^2+^ crystal structure (1vfp), the γ-phosphate of the ATP analog, AMPPCP, establishes hydrogen bonds with these two residues, T353 and T625, whereas, in the crystal structure of SERCA1a.Mg^2+^:SLN (3w5a) the γ-phosphate of TNPAMP is far from them. This difference could be attributed to the nature of the ATP analogs in the two structures (1vfp and 3w5a), since TNPAMP is not as similar to ATP as AMPPCP. However, in both our structures, where TNPAMP was replaced with ATP (see Methods for details), this different position of ATP is still observed after energy minimizations. Indeed, the ATP γ-phosphate stays far from T353 and T625 in the presence of SLN, whereas in the absence of SLN, it comes close and establishes hydrogen bonds with the hydroxyl groups of the two threonines, as in the E1.2Ca^2+^ crystal structure. This shows that the distance between the ATP γ-phosphate and the two threonines is influenced by the presence of SLN.

However, ATP is in the part of the protein that is accessible to solvent and is therefore surrounded by the shell of water and ions that we considered in our calculations. This shell is the same for the two systems E1.Mg^2+^ and E1.Mg^2+^:SLN. To confirm our results, a second calculation in the presence of SLN was performed, using a different water/ions shell that was randomly created (see Methods). All the results were similar to those presented throughout this article, showing their robustness and independence from the water/ion layer.

Analyses of both energy-minimized structures, with and without SLN, showed that the region around one of the two threonines (T353) and that consists of residues 352 to 357, is unstructured in the presence of SLN. This region is called here the P-N linker because it links the P- and N-domains (Figure 7C-E). This P-N linker adopts a π−helix structure in the absence of SLN, by establishing a hydrogen bond between the C=O group of K352 and the N-H group of T357. Remarkably, this region almost encompasses the entire autophosphorylation site i.e. the ^351^DKTG^354^ motif. Due to the structuration of the P-N linker, which involves the backbone of K352, a small torsion of the second (residues 347 to 352) and third (residues 620 to 625) β−strands of the central β−sheet of the P-domain takes place. This is because K352, which starts the P-N linker, also ends the second β−strand (Figure 7C-E). Since the third β−strand ends with T625, this torsion brings the N-H group of T625 to establish a hydrogen bond with the ATP γ-phosphate. In addition, due to the structuration of the P-N linker, both N-H and O-H groups of T353 establish H-bonds with the same oxygen atom of the ATP γ-phosphate. These interactions add up to the one that exists, in both the presence and absence of SLN, between the C=O group of T353 and the Mg^2+^ chelated to ATP. Therefore, starting from the same initial structure, 3w5a, energy minimizations in the absence of SLN brought the protein to a conformation which stabilizes the ATP γ-phosphate in an intermediate position that precedes autophosphorylation, whereas, this was not the case in the presence of SLN. So the question is how SLN impedes the structuration of the P-N linker while it is 35 Å away.

### 7-…due to the straightening of TM6 and TM5

In both energy-minimized structures, E1.Mg^2+^:SLN and E1.Mg^2+^, the P-N linker is in close contact with F740, which is the N-terminus of helix TM5 (Figure 7C-E). The N-terminal half of this long helix (residues 740 to 759) is cytoplasmic, whereas its C-terminal half (residues 760 to 780) is embedded in the membrane and in contact with TM6 (residues 789 to 810). The curvature of the cytoplasmic half of TM5 changes (Figure S4 in Supplemental Information); it straightens in the presence of SLN due to the relative straightening of the 3-10 helix of TM6, which takes place because of the hydrogen bond of G801 and T805 of this helix with N11 of SLN, as described above in subsection 5. Indeed, the straightening of TM6 results in the reorientation of the neighboring L807, which comes closer to Q759 of TM5 and pushes it, promoting TM5 straightening. The produced steric hindrance explains the decrease of the fluctuation of residues Q759, L807 and G808 in the presence of SLN as presented above in subsection 4 and Figure 5A. Mutations of these residues show their importance for SERCA1a activity. Indeed, the mutant Q759A presents a lower turn-over rate (74%) than the wild type (WT), which is accompanied by phosphorylation slowdown (t_1/2_ is 0.26 s^-1^ for WT and 0.43 s^-1^ for Q759A) (52). Additionally, mutant L807A has a significantly reduced affinity for calcium (Ca_1/2_ drops from 0.35 µM for WT to about 5 µM for mutant) but with a moderate effect on the calcium transport rate. Mutant G808A is much slower (only 50% of the calcium transport rate of the WT), although with only a marginal effect on calcium binding (Ca_1/2_ of about 0.3 µM for the mutant (62)).

The difference of curvature of TM5 has an effect on the relative position of the P-N linker: in the presence of SLN, the C=O group of T355, which is located in the middle of the linker, is equidistant from the two N-terminal residues of TM5, N739 and F740 (the distance of the carbonyl O of T355 from the C_α_ aliphatic hydrogen of N739, O−H_α_ = 2.5 Å and from the hydrogen of the amide group of F740 O-HN = 2.5 Å), whereas in its absence, this carbonyl group is slightly closer to F740 (distance O-HN = 2.4 Å) but significantly farther from N739 (distance O−H_α_ = 3.3 Å). Consequently, in the presence of SLN, the P-N linker is more constrained, as it is stuck on the top of TM5 equidistantly from N739 and F740, and therefore, its structuration is impeded. Site-directed mutagenesis has indeed demonstrated that mutations of all the residues of the P-N linker, except G354A, result in a slow enzyme phosphorylation turnover, showing the importance of this linker in the phosphorylation process (61). The N-terminal part of TM5 has also a crucial role in mediating communication between the calcium binding sites and the catalytic domain (52), which was shown even before resolving the first crystal structures (63). Replacement of F740 with a leucine, a residue almost as bulky as phenylalanine, moderately affects the enzyme turnover rate (83% of the WT), whereas this rate drops down to less than 10% when F740 is replaced with an alanine (52).

## Discussion

### The metal binding site

Comparison of the various crystal structures of the E1.2Ca^2+^ state, like 1vfp and 3ar2, shows that the Ca^2+^ binding sites I and II are well defined, with similar positions of the two Ca^2+^ ions and their chelating residues (for 1vfp, see Figure 6A). This is not the case for the crystal structures obtained in presence of Mg^2+^, whether SLN is present or not. Indeed, the Mg^2+^ binding site I’, is not well defined because there are doubts about the position of Mg^2+^ and its chelating residues (Figure 6B, D, E), because of a too poor resolution in that region, as mentioned by the authors (13). In the crystal structures 3w5a (SERCA1a.Mg^2+^:SLN) and 3w5b (SERCA1a.Mg^2+^ in the absence of SLN), the cation is placed near N768 and far from D800, whereas in 4h1w (SERCA1a.2Mg^2+^:SLN), the cation in site I’ is placed near D800 and far from N768. In the latter structure a second cation is trapped at the entrance of the binding sites between E309 and V304, “stabilizing an open structure” (15). Considering the two energy-minimized structures generated here from 3w5a, in the absence of SLN, the Mg^2+^ ion is chelated by both N768 and D800, whereas in its presence, D800 cannot come close enough to chelate Mg^2+^ because of the hydrogen bonds between the neighboring G801, T805 on the one hand and SLN-N11 on the other hand (Figure 7B). We assume that this would also be the case in the presence of Ca^2+^, or maybe, since Ca^2+^ is bigger than Mg^2+^, D800 would possibly be able to chelate only one Ca^2+^ but not two as it does in the absence of SLN. This would, in any case, slightly reduce the affinity for calcium in the presence of SLN, which is in good agreement with the effect of sarcolipin on the Ca_1/2_ that drops from 0.35 ± 0.02 µM to 0.51 ± 0.02 µM as described earlier (17). Remarkably, the mutation of SLN-N11 to alanine restores almost completely the calcium affinity with a Ca_1/2_ 0.39 ± 0.03 µM. Additionally, by its proximity to G801 and T805, which directly interact with SLN, and also its proximity to TM5, D800 seems to play a role, not only in the binding of the cation, but also in the transmission of information from the Ca^2+^ binding sites to the phosphorylation site. The observation of the experimental effects of mutations D800N and D800E, in the absence of SLN, shows that both the calcium transport and the phosphorylation turnover are 2 fold slower (44, 45, 48, 53, 54).

At the entrance of the Ca^2+^ binding sites, the distance between the C_α_ atoms of V304 and E309 in E1.Mg^2+^ is slightly smaller than in E1.Mg^2+^:SLN. Such a difference is also measured between these C_α_ distances within the PDB structures, 3w5a and 3w5b. However, these two residues are substantially more flexible in the absence of SLN, probably due to the absence of strain exerted by the peptide on the C-terminus of helix M4S4. This flexibility is necessary, at least for E309, to chelate the cation. The lower flexibility of these residues in the presence of SLN may also result in a slight reduction of Ca^2+^ affinity.

### The phosphorylation site

The two energy-minimized structures E1.Mg^2+^:SLN and E1.Mg^2+^ are similar with an RMSD over all C_α_ atoms around 1 Å. Nevertheless, some important local differences were highlighted in the results, notably in the P-N linker, in the TM5 and TM6 helices and in the position of ATP. Indeed, in the presence of SLN, the P-N linker is unstructured as in the E2-state crystal structures (3w5c, 2dqs) and TM5 and TM6 helices are straightened (Figure S4A in Supplemental Information). ATP is in a relatively high, deep position into the N-domain − with a distance between the top ATP atom (N1) and its closest residue M494 (atom S) of 3.93 Å −, which does not allow it to establish H-bonds with T353 and T625 (for the distances between the ATP γ−phosphate and these residues, refer to the legend of Figure 7C, D). In the absence of SLN, the P-N linker adopts a π−helix structure, helix TM5 is slightly more curved and TM6 slightly more kinked than in the presence of SLN. Therefore, ATP can come in a lower and shallower position relative to the N-domain (the distance ATP:N1−M494:S = 5.15 Å), allowing it to establish H-bonds with both T353 and T625.

These differences are also observed, at least qualitatively, between the two crystal structures SERCA1a.Mg^2+^:SLN (3w5a) and SERCA1a.Mg^2+^ (3w5b), although these structures are overall similar (RMSD ≈ 1 Å). Indeed, in 3w5b (SERCA1a.Mg^2+^), the P-N linker is structured, helix TM5 slightly more curved and TM6 slightly more kinked than in 3w5a (SERCA1a.Mg^2+^:SLN) (Figure S4B in Supplemental Information). However, TNPAMP, the ATP analog in these crystal structures, is not close to T353 and T625 in 3w5b (SERCA1a.Mg^2+^), like in the E1.Mg^2+^ energy-minimized structure. This is probably due to strong electrostatic interactions (or H-bonds although they are not observed as such) between the nitro groups of TNPAMP (which are not present in ATP) and surrounding arginines and lysines (K515, R560 and R678) of the N-domain. Therefore, in the crystal structure 3w5b (SERCA1a.Mg^2+^), the position of TNPAMP is not relevant for the protein activity. On the other hand, the energy-minimized structure E1.Mg^2+^ was compared to 1vfp and 3ar2, the two crystal structures of E1.2Ca^2+^, comprising the SERCA1a disulfide bridge and an ATP analog, AMPPCP, more similar to ATP. The closer position observed in E1.Mg^2+^ of ATP with T353 and T625 residues was also observed in 1vfp and 3ar2. In the latter structures, the γ−phosphate establishes H-bonds with the hydroxyl groups of both T353 and T625 as in E1.Mg^2+^. Remarkably, in these structures, the P-N linker adopts a π−helix structure accompanied by the curvature of TM5 and the kink of TM6, as in 3w5b (SERCA1a.Mg^2+^) and our energy-minimized E1.Mg^2+^ structure. These local modifications around the phosphorylation site, may slightly slow down the ATP hydrolysis rate as observed in (18, 64–72).

### Comparison of the effects of sarcolipin and phospholamban on SERCA1a

The structure of SERCA1a in the presence of another regulatory peptide, phospholamban (PLN), is also known (73). PLN is mainly expressed in cardiac and smooth muscle cells (74). It is a peptide of 52 residues composed of a cytosolic N-terminal part of 23 residues and a transmembrane C-terminal part of 29 residues. PLN binds to SERCA1a in a way similar to SLN and it has similar effects on the protein activity (75), although the sequence alignment of the TM part of PLN and SLN shows that only two adjacent residues are fully conserved, namely N11 and F12 of rabbit SLN. The alignment was done with the SEAVIEW program (76, 77) using all non-redundant sequences from various organisms given in UniProt (https://www.uniprot.org), i.e., 108 organisms for PLN and 110 for SLN. Interestingly, the conserved residue N11 contributes to the straightening of SERCA1a TM6 by establishing an H-bond with SERCA1a-T805, which is also fully conserved in 116 non-redundant sequences. Comparison of the structure of SERCA1a-PLN (4kyt) with those of SERCA1a-SLN (3w5a and 4h1w) shows that they are similar, despite the absence of cations in the Ca^2+^-binding sites in the structure with PLN. However, SERCA1a-PLN is closer to 3w5a (RMSD = 1.6 Å on all C_α_ atoms) than to 4h1w (RMSD = 2.6 Å). In the structure of SERCA1a-PLN (4kyt), the P-N linker is unstructured and helices TM5 and TM6 straightened like in all the crystal structures of SERCA1a-SLN (3w5a, 4h1w) and in the energy-minimized structure E1.Mg^2+^:SLN. This observation suggests that SLN and PLN follow a similar mechanism for the inhibition of SERCA1a, especially that this inhibition is lost when SERCA1a-T805 is mutated to alanine (62, 78, 79).

### Deciphering the propensity for the transition toward E2

The results presented above revealed that, in the presence of SLN, the motions of SERCA1a toward the E2 state are possible, whereas those toward the E1.2Ca^2+^ state are penalized, although E1.Mg^2+^:SLN is already in an E1-like state. Conversely, in the absence of SLN, although the structure is similar to that in presence of SLN, the motions toward E2 are penalized, whereas the transition toward the E1.2Ca^2+^ state is now more favorable. This tends to confirm that the E1.Mg^2+^:SLN state is an intermediate conformation between E2 and E1.2Ca^2+^, as proposed by Toyoshima *et al.* (13) and Winther *et al.* (15), and that SLN impedes the motions that bring the structure close to E1.2Ca^2+^. Analyses of the motions along the modes and of the structural differences between E1.Mg^2+^:SLN and E1.Mg^2+^ provide us with an explanation to these observations. Indeed, to go toward E2, the P-domain undertakes a rotation around helix TM5 and entails the rotation of the N-domain in the same direction and of the A-domain in the opposite direction, like a cogwheel. In the presence of SLN, the peptide pulls the 3-10 helix part of TM6 a little away from TM5 (except for L807 whose sidechain then straightens and comes closer), leaving enough room for TM5 to straighten up and to adapt to the P-domain rotation, which makes this rotation, and therefore the transition toward E2, comfortable. In the absence of SLN, TM6 leans on TM5, modifying the curvature of the helix and increasing its interactions with the P-domain, which makes the independent rotation of the domain, and therefore the transition toward E2, more difficult.

### Deciphering the propensity for the transition toward E1.2Ca^2+^

For the transition toward E1.2Ca^2+^, the N-domain undertakes, with the A-domain, a closing movement, similar to the hinge-bending motion. This movement may be described as a rotation of the N-domain around the axis of the P-N linker. In the presence of SLN, concomitantly to this rotation, TM5 undertakes an upward movement, whereas this helix is rather straight, with almost all its expected *i, i*+4 backbone H-bonds present. So TM5 behaves as a rigid body along its axis, and during its upward movement, it squeezes the unstructured P-N linker, which sits on the top of it, pushing it toward the N-domain. This movement presents an important stress and results in an anticorrelation between the N-domain and TM5, as observed in the heatmap of mode 41 of E1.Mg^2+^:SLN (Figure 4D). In addition, because ATP is positioned a little deeper into the N-domain, compared to its position in the absence of SLN, the N-domain encounters more difficulty during its rotation movement. All this together impedes the transition toward E1.2Ca^2+^ in the presence of SLN.

In the absence of the peptide, ATP is in a shallower position, releasing the N-domain rotation. In addition, the P-N linker is structured and slightly moved away from the top of TM5, which allows a little sliding of TM5 around the linker. TM5 does not undertake an upward movement similar to that in the presence of SLN, hence the absence of its anticorrelations with the N-domain (Figure 4C), but its sliding around the linker is accompanied by a slight compression of the helix along its axis. However, since TM5 is rather curved, with a few number of its *i, i*+4 backbone H-bonds disrupted, it is less rigid and its slight compression is softens, which makes the transition of the protein toward E1.2Ca^2+^ possible in the absence of SLN.

### The central role of the P-domain

The transition toward either E2 or E1.2Ca^2+^ proved to be governed by either the motion of the P-domain or around the P-N linker, which is usually considered as part of the P-domain. These motions are determined by the conformation of helix TM5, which, in its turn, is affected by the structural modifications of TM6 due to the presence of SLN. The importance of the P-domain in the state transition of SERCA1a was hypothesized by Møller *et al.* (5) based on structural analyses of the protein, in addition to the conservation of this domain. Indeed, the P-domain is the most highly conserved part of SERCA, and this is true for P-type ATPases in general (80).

## Conclusion

Based on Normal Modes calculations and Energy Minimizations, we may summarize the mechanism of action of SLN on SERCA1a as follows. When SLN binds to SERCA1a, it establishes hydrogen bonds between its N11 residue on the one hand and SERCA1a T805 and G801 on the other hand, which induces a pulling of the 3-10 helix part of TM6 toward SLN and away from TM5. This movement has effects on both the Ca^2+^ binding sites and the phosphorylation site. For the former, a constraint is then introduced on D800, which is located in TM6, preventing it from chelating the metal. Concomitantly, the two gating residues at the entrance of the binding sites, V304 and E309, become less flexible, making the entrance of the metal more difficult. This decreases the affinity of SERCA1a for Ca^2+^, although the structure is in an E1-like state. The second effect is on the phosphorylation site, more than 35 Å away from SLN. This effect is mediated by helix TM5, which straightens when TM6 is pulled away from it by SLN. The straightening up of TM5 destructures the P-N linker that sits above it, hampering ATP from coming close to D351 (the residue to phosphorylate), and therefore decreasing the phosphorylation rate. Besides, the presence of SLN hinders the transition toward the E1.2Ca^2+^ state because of the position of ATP, deeper in the N-domain, and of the destructuration of the P-N linker, which impede the sliding of the N-domain around this linker for a hinge-bending-like motion. Conversely, the transition toward E2 is facilitated by the presence of SLN, because, by moving TM6 a little away from TM5, SLN contributes to reducing steric hindrance on TM5, enabling it to adapt to the P-domain rotation, necessary for this transition. Finally, this mechanism can be generalized to other inhibitory peptides, such as PLN, that bind to SERCA1a in a manner similar to that of SLN.

## Acknowledgments

The calculations were performed on equipment funded by the French program “Investissement d’Avenir – Institut Carnot” managed by the National Research Agency (ANR-11-CARN-008-01). T.B. benefited from a PhD fellowship from Saclay University (ED568-Biosigne).

## Author Contributions

TB did the calculations under the supervision of LM and VB; TB and LM analyzed the results and wrote the first draft of the article; EQ adapted the method DIMB for GPU and tested it on small proteins; VB, CM and NJ provided the subject, compared the simulation results to experimental results and participated in the manuscript writing.

